## Supplementary material for "The landscape of coadaptation in *Vibrio parahaemolyticus*": Suppmentary Figures

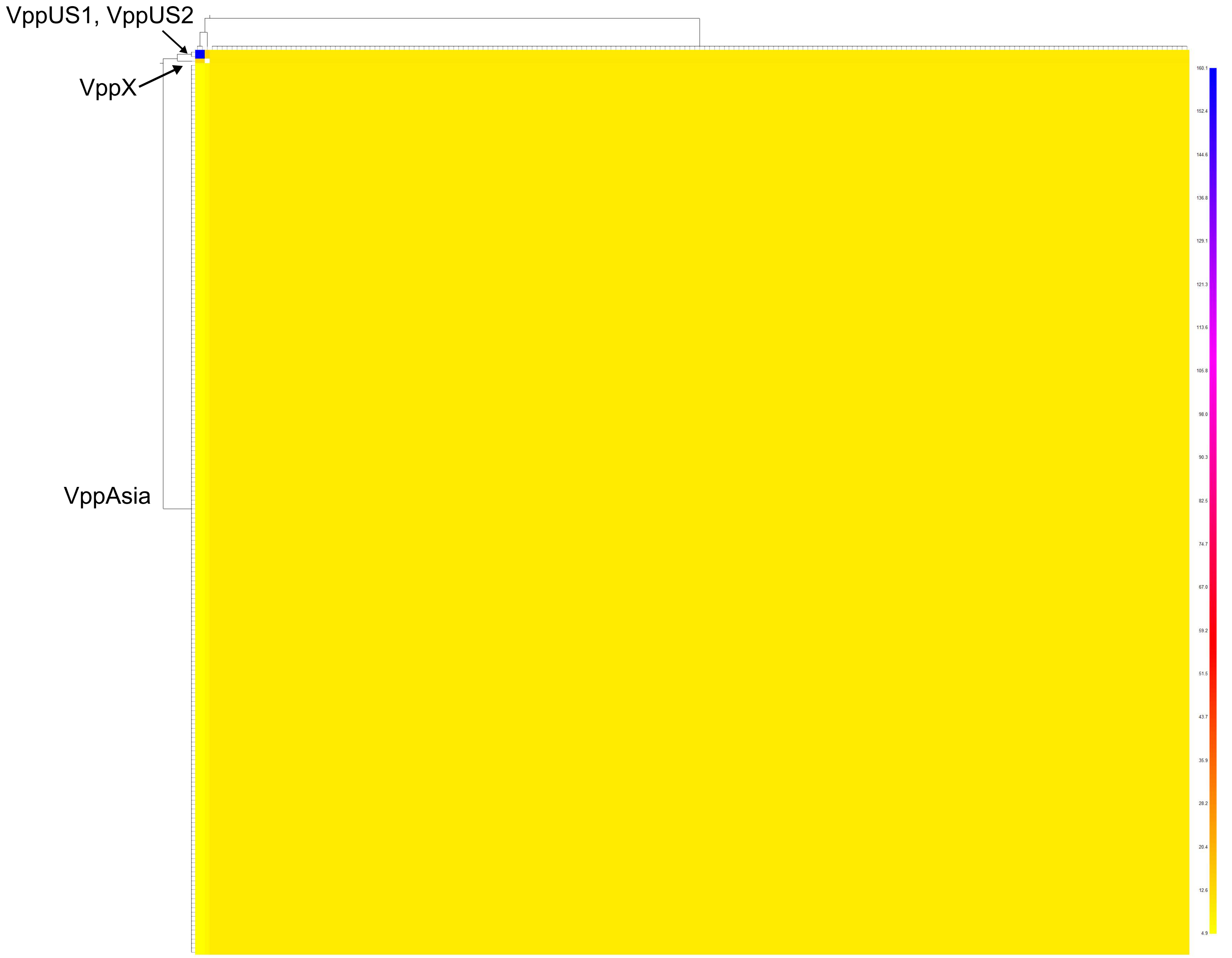


**Figure S1. fineSTRUCTURE showing no clonal frame in 198 VppAsia isolates.** Coancestry matrix of 201 strains, including 198 VppAsia isolates and three isolates respectively from VppX, VppUS1 and VppUS2. The color of each cell indicates the expected chunks numbers imported from a donor (column) to a recipient (row).

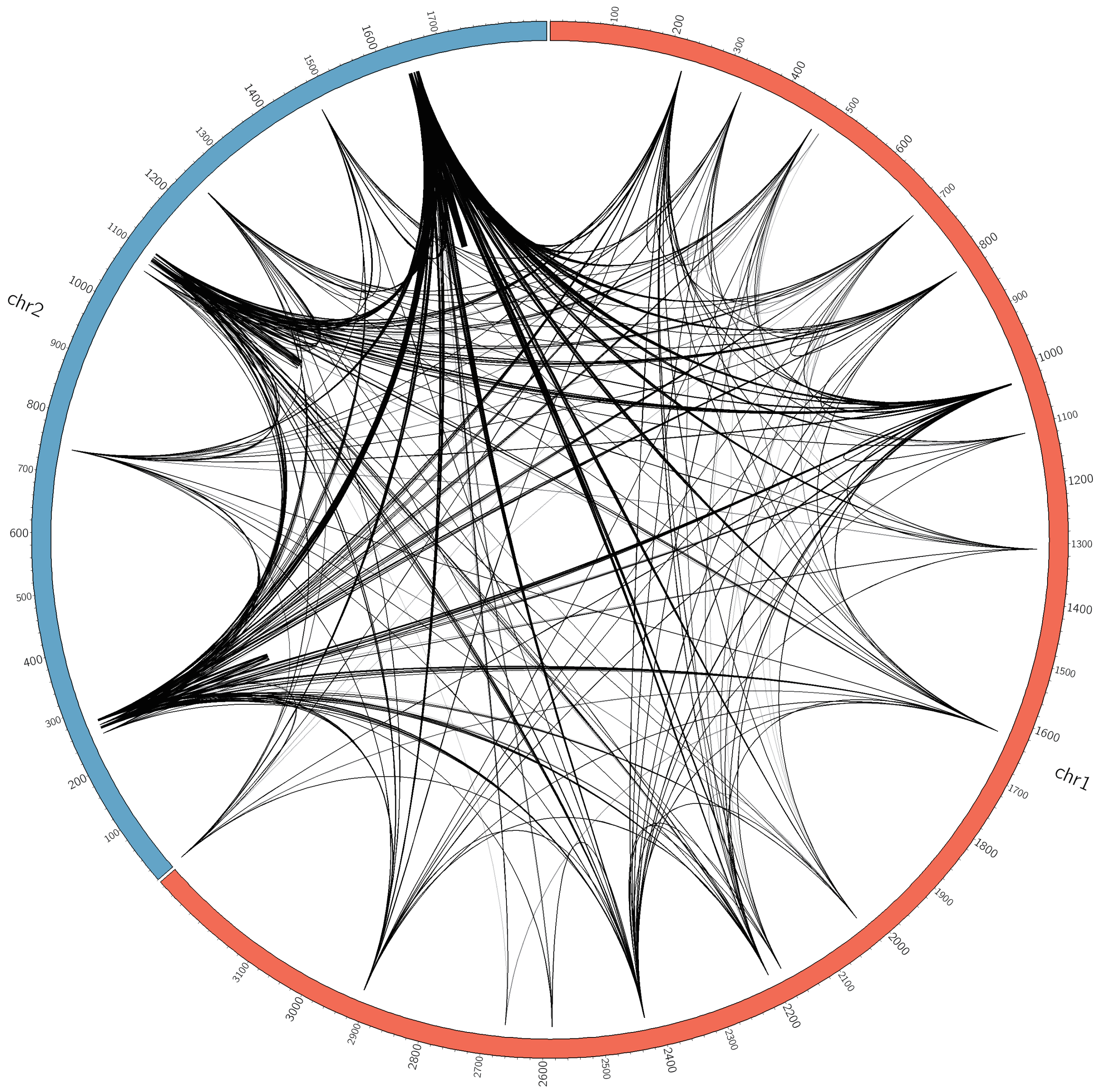


**Figure S2. EG1a interactions that only detected by Fisher exact test.**

**FigureS3. Clustering of 469 strains based on variations of IG2-90.** Hierarchical clustering of strains (column) based on coadaptation loci (row) of all IGs, except for IG1, 2, 3, 14, and 42 that had been showed in figures of the main text. Color scheme is the same as in Figure 3.

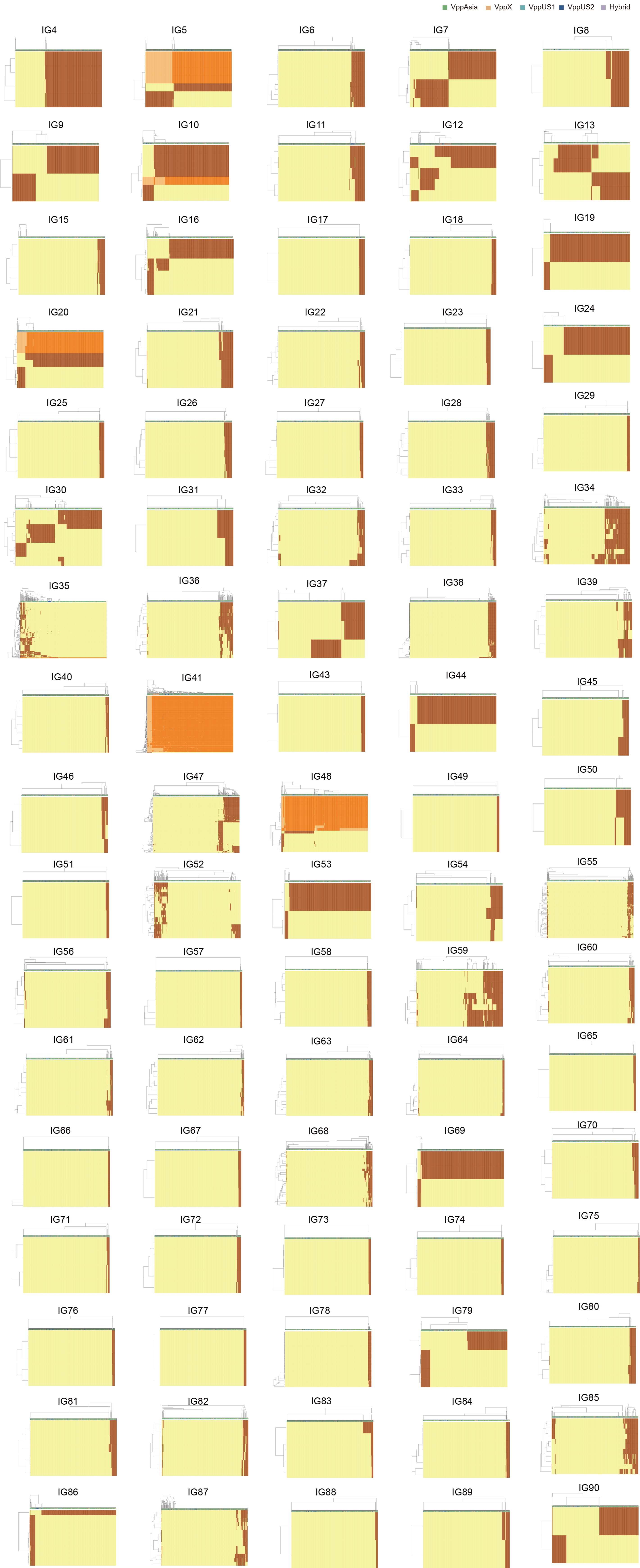

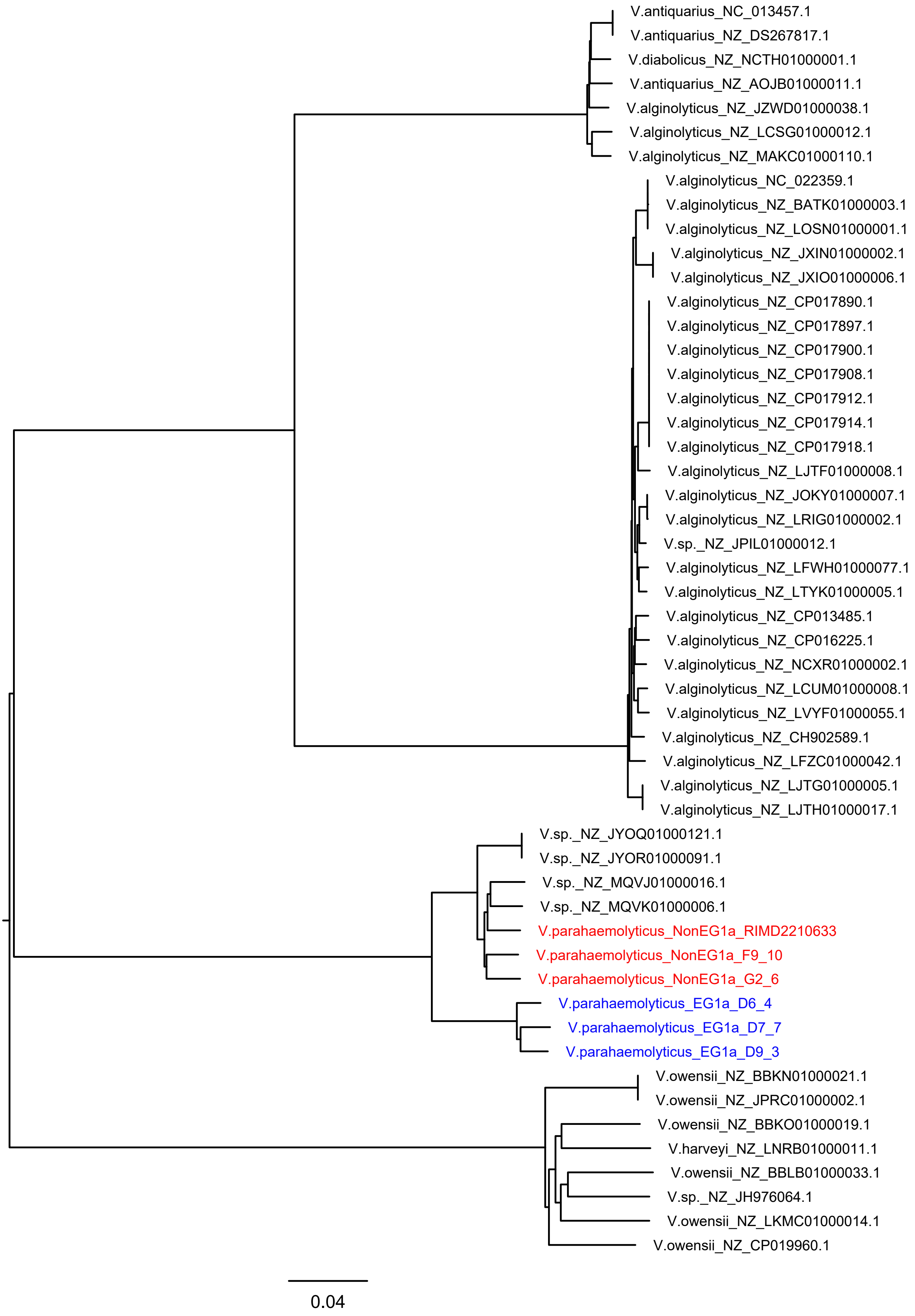


**Figure S4. NJ tree of lateral flagellar gene cluster region (VPA1538-1557) in *Vibrio* genus.** Three randomly selected strains of EG1a and NonEG1a are used and marked in blue and red, respectively.

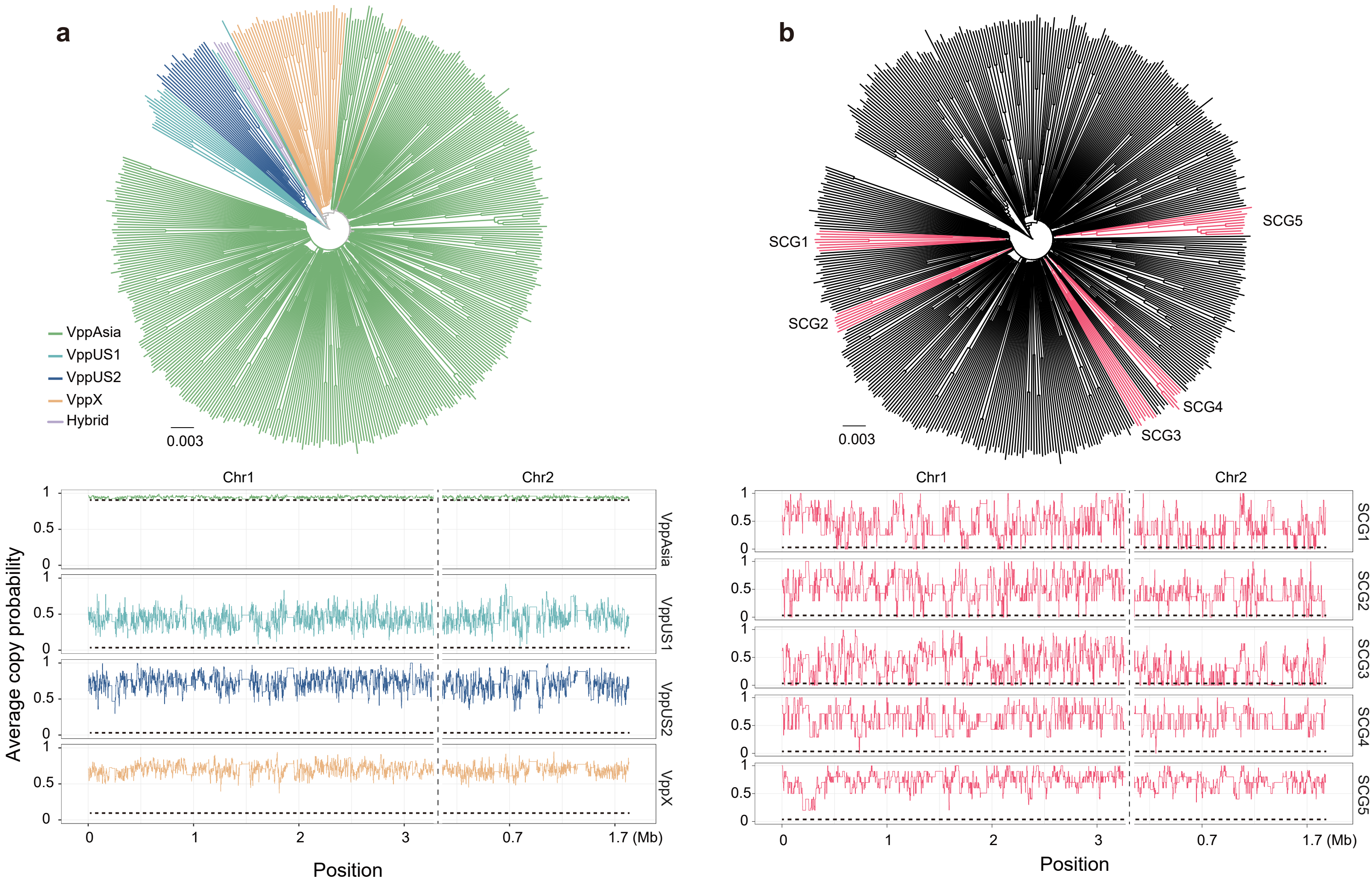


**Figure S5. NJ trees (top) and average copy probability value distributions (bottom) of geographical populations (a) and semi-clonal group (SCG) strains (b) of *V. parahaemolyticus*.** Branch colors of the trees indicate populations (a) and SCGs (b), respectively. Horizontal dotted line indicates the expected average copy probability value. Different chromosomes are separated by a vertical dotted line.
